# Dynamics of mixed-ploidy populations under demographic and environmental stochasticities

**DOI:** 10.1101/2023.03.29.534764

**Authors:** Michelle L. Gaynor, Nicholas Kortessis, Douglas E. Soltis, Pamela S. Soltis, José Miguel Ponciano

## Abstract

The theoretical population dynamics of autopolyploids – organisms with more than two genome copies of a single ancestral species – and their diploid progenitors have been extensively studied. The acquisition of multiple genome copies, being in essence a stochastic process, strongly suggests a probabilistic approach to examine the long-term dynamics of a population with multiple cytotypes. Yet, our current understanding of empirical evidence on the dynamics of autopolyploid populations has not incorporated stochastic population dynamics. To investigate the factors contributing to the probability and stability of coexisting cytotypes, we designed a new population dynamics model with demographic and environmental stochasticities to simulate the formation, establishment, and persistence of diploids, triploids, and autotetraploids over time when gene flow is allowed among cytotypes. Contrary to previous research, increased selfing rates and pronounced reproductive isolation stabilized the long-run coexistence of multiple cytotypes. In stressful environments, these dynamics become much more complex, and our stochastic modeling approach helped reveal the resulting intricacies that give tetraploids competitive advantage over their diploid progenitors. Our work is fundamental to a better understanding of the dynamics of coexistence of multiple cytotypes and is a necessary step for further work modeling the dynamics between an autopolyploid and its diploid progenitor.

## Introduction

Polyploid formation, or whole-genome duplication (WGD), is observed across the tree of life (Conant 2020; Gregory and Mabel 2005; Initiative 2019; Li et al. 2018; Schmid et al. 2015; Spoelhof et al. 2019) and is in essence a stochastic process. Yet, current literature is marked by a paucity of rigorous explorations into the joint effects of demographic variability and environmental stochasticity in polyploid persistence. Changes in extinction or persistence probabilities in nature are best understood using both deterministic and probabilistic tools (Cohen 2004; Cushing et al. 2003). Incorporating stochasticity is thus expected to markedly improve our understanding of complex dynamics in naturally occurring populations. To investigate coexistence among organisms that differ in the number of copies of their genome (cytotypes), we designed a new matrix population dynamics model with demographic and environmental stochasticities to simulate the formation, establishment, and persistence of diploids, triploids, and tetraploids over time when gene flow is allowed.

WGD is an important evolutionary force among angiosperms (Leitch and Bennett 1997; Ramsey and Ramsey 2014; Soltis et al. 2009). On one end of the spectrum, genome duplication may involve one individual or occur following crossing between different individuals of the same species, known as autopolyploidy; on the other end of the spectrum, genome duplication may occur in conjunction with hybridization between individuals of two separate species, known as allopolyploidy. Autopolyploids are mostly formed through the union of unreduced gametes or a multi-generational process involving a triploid bridge (Harlan and deWet 1975). An alternative pathway is through somatic doubling of a cell line involved in flower production (Schoen and Schultz 2019); however, this is considered rare in nature (reviewed in Soltis et al. 2016).

After formation, the fate (*i.e.*, establishment or extinction) of the newly formed autotetraploid is stochastic; the chance of establishment or extinction is dependent on its vigor and competitive ability (Harlan and deWet 1975). The establishment of a newly formed autopolyploid is limited by minority cytotype exclusion (MCE), the process wherein the rare cytotype has limited mate availability and therefore faces negative selection pressure (Felber 1991; Fowler and Levin 1984; Levin 1975; Rausch and Morgan 2005). MCE is an example of positive frequency-dependent selection and represents an ecological Allee effect (Dennis 1989).

The maintenance of multiple cytotypes (cytotype coexistence) requires mechanisms that overcome MCE, and MCE has been the focus of theoretical models for the last five decades (Baack 2005; Burton and Husband 2001; Felber 1991; Felber and Bever 1997; Fowler and Levin 1984; Husband 2004; Levin 1975; Oswald and Nuismer 2011; Rausch and Morgan 2005; Rodriguez 1996; Spoelhof et al. 2020; Suda and Herben 2013; Van Drunen and Friedman 2022; Yamauchi et al. 2004). Early deterministic models investigated MCE for diploids and autotetraploids in relation to self-compatibility (Levin 1975), viability and fertility (Felber 1991), and with explicit competition as defined by Lokta-Volterra models (Fowler and Levin 1984; Rodriguez 1996). These models found that under certain conditions, for example, niche separation between cytotypes, successful long-term establishment of autotetraploids is possible alongside diploids despite MCE. Taken together, previous theoretical models conclude that establishment of autotetraploids can be aided by recurrent formation, reproductive assurance, and reproductive separation.

Most mixed-cytotype autopolyploid systems examined to date appear to have experienced multiple origins in their history (*i.e.*, recurrent formations), rather than a single origin (Husband and Sabara 2003; Levin 2011; Parisod and Besnard 2007; Ramsey et al. 2008; Segraves and Thompson 1999; Servick et al. 2015; Těšitelová et al. 2013). The probability of success of the newly formed autotetraploid (*i.e.*, a neoautotetraploid) increases with recurrent formation. However, an autopolyploid could form a single time from a diploid parent and become successfully established, as is likely in *Arabidoposis arenosa* (Arnold et al. 2015) and possibly *Tolmiea menziesii* (Soltis et al. 1989; Visger et al. 2016). This pattern suggests that a neoautotetraploid has a low probability of persistence, but this may be overcome by repetitive WGD events.

Recurrent formation can occur through two main paths. First, an autopolyploid (here, an autotetraploid) may form from the union of two unreduced gametes by either a single diploid individual or by two individuals of the diploid species; formation through this path can be represented as a joint probability. Second, an autotetraploid may form through a triploid bridge, in which a triploid intermediary forms via the union of a reduced gamete (1*n*) and an unreduced gamete (2*n*) from the diploid progenitor (again, either a single individual or two individuals). This triploid intermediate can generate gametes that are 1*n* (average 3%), 2*n* (average 2%), 3*n* (average 5.2%), or aneuploid (average 89.8%) (values from Ramsey and Schemske 1998). Triploid gametes within a population can then unite with reduced or unreduced gametes produced by diploids (and autotetraploids); in rare instances, these unions will result in a viable combination (with the triploid being a triploid bridge); however, these combinations are often unsuccessful, incompatible, and sterile (a triploid block) (Kö hler et al. 2010). Therefore, probability of a successful autotetraploid offspring forming due to a triploid bridge is weighted by the proportion of viable gametes formed and the probability of this gamete to unite with a compatible gamete. Additionally, triploids may promote the maintenance of diploids (Felber and Bever 1997), aid in establishment of autotetraploids (Burton and Husband 2001), and likely aid in the persistence of mixed-cytotype populations (Husband 2004). However, the impact of triploids may be modulated by the selfing frequency and ability of all cytotypes within a population (Yamauchi et al. 2004). Despite triploids influencing the probability of coexistence of diploids and autotetraploids, most theoretical models have omitted triploids. Although triploids may have a critical role in establishment and persistence, aiding in both recurrent formation and later gene flow, triploid trajectory is not a focus in our current efforts as triploids are expected to have low fertility (Kö hler et al. 2010; Ramsey and Schemske 1998) and thus exist at low frequencies.

Models predicting success (*i.e.*, establishment, followed by persistence) of a neoautopolyploid have found that it is increased by reproductive assurance through selfing (Baack 2005; Oswald and Nuismer 2011; Rausch and Morgan 2005; Rodriguez 1996) or clonality (Chrtek et al. 2017; Van Drunen and Friedman 2022). In naturally occurring populations, autopolyploids have been observed to experience shifts in mating system, often to facultative reproductive systems with both asexual and sexual methods of reproduction (Barringer 2007; Husband et al. 2008). The breakdown of self-incompatibility and the production of offspring via selfing in neoautote-traploids also facilitates establishment, often as colonization of new locations such as islands; self-compatible individuals are expected to be more successful at colonization and establishment than self-incompatible individuals due to the ability of the former to produce offspring without mates (Baker 1967).

Reproductive isolation through pre-zygotic mating barriers can limit secondary contact and interactions between multiple co-occurring cytotypes, thereby promoting establishment and persistence. Theoretical studies have suggested that the success of the neoautopolyploid is weighted by its interactions with its diploid progenitor and that decreased interactions between the cytotypes promote establishment (Baack 2005; Griswold 2021; Rodriguez 1996; Spoelhof et al. 2020; Van Drunen and Friedman 2022). Reproductive isolation and cytotype separation can be achieved through pre-zygotic barriers such as trait divergence (*e.g.*, flowering time and pollinator preference) or spatial segregation. Shifts in flowering time may limit reproduction between cytotypes; for example, the flowering time of autotetraploid *Anacamptis pyramidalis* (Orchidaceae) is shifted from that of diploids, thus reducing competition for pollinators and allowing similar levels of reproductive success for each cytotype (Pegoraro et al. 2019).

Microgeographic organization in naturally occurring populations of mixed cytoypes has rarelybeen investigated; however, theoretical models have repeatedly suggested that spatial segregation is essential for neoautotetraploid persistence (Baack 2005; Griswold 2021; Kauai et al. 2023; Spoelhof et al. 2020; Van Drunen and Friedman 2022). Despite the theoretical expectation of spatial divergence, mixed cytotype populations lacking spatial segregation have been observed in many taxa (as reviewed by Kolář et al. 2017). It is unknown if these observations of mixed cytotypes are merely ephemeral ‘snapshots’ (Levin 1975; Mráz et al. 2022); however, persistence of an autotetraploid in natural populations is difficult to monitor, and temporal observations of naturally occurring mixed-ploidy populations are few: for example, a stable perennial system (*Vicia cracca* (Fabaceae); Trávńıček et al. 2010) and two systems (annual and mixed life history) which are persistent despite fluctuations in the relative frequency of cytotypes (*Centaurea stoebe* (Asteraceae) and *Tripleurospermum inodorum* (Asteraceae); Mráz et al. 2022; C^̌^ ertner et al. 2019).

In both theoretical and empirical studies, long-term persistence (*i.e.*, success) of coexisting autotetraploids is difficult to define. Stable equilibria have been investigated in numerous mixed-cytotype models (Burton and Husband 2001; Felber 1991; Felber and Bever 1997; Fowler and Levin 1984; Levin 1975); however, persistence in stochastic conditions has not been explored. Here, we explore persistence with a simulation-based approach that incorporates the effects of both demographic and environmental stochasticities into population dynamics, thus incorporating chance variation in birth and death processes as well as random temporal variation in the per-capita growth rates (Lewontin and Cohen, 1969).

Because coexistence with incomplete reproductive isolation has seldomly been modeled (Ir-win and Schluter 2022), here we attempt to better understand the dynamics of coexistence of multiple cytotypes under various levels of reproductive isolation through established probabilistic tools in evolutionary ecology and population dynamics. Although there are well-established methods and coexistence metrics that rely on invader growth rates (*e.g.*, Chesson 2018; Schreiber et al. 2011), they are applicable when species are reproductively isolated and do not apply to the cases we study here. Importantly, in a typical stochastic multi-species ecological model with reproductive isolation, extinction occurs when the abundance of a particular taxon reaches zero (an abundance of zero is then defined as an “absorbing state”, Allen 2010); however, key idiosyncrasies of polyploid populations render traditional tools and concepts in population dynamics inadequate. In particular, any given species or type can re-appear after reaching an abundance of zero in the scenario we consider here: for example, via recurrent formation of a tetraploid or recurrent formation of a triploid via crossing between a diploid and a tetraploid. Thus, absorbing states no longer exist, and classic extinction results of stochastic population dynamics do not apply (except in the limiting case of all cytotypes going to zero). To our knowledge, our study is the first to approach these theoretical and conceptual needs through modern probabilistic population dynamics modeling.

## Methods

To better understand formation, establishment, and persistence of diploids, triploids, and tetraploids (*i.e.*, autotetraploids), we formulated a stochastic, stage-structured matrix population dynamics model (Caswell 2000) of a mixed-cytotype perennial plant population with overlapping generations. The model incorporates both demographic and environmental stochasticities (Ovaskainen and Meerson 2010). We assumed a demographically closed population (*i.e.*, no emigration or immigration from additional sources) such that the demographic processes defining population changes in both size and composition from one time step to the next are reproduction, survival, and maturation to reproductive maturity (Fig. 1). Although this model is complex, through extensive simulations summarized below, we can nonetheless characterize the effects of reproductive isolation, founding population size, and variation in the environment on: (1) the probability that diploids and tetraploids coexist and (2) the resulting stability of coexistence.

**Figure 1:**
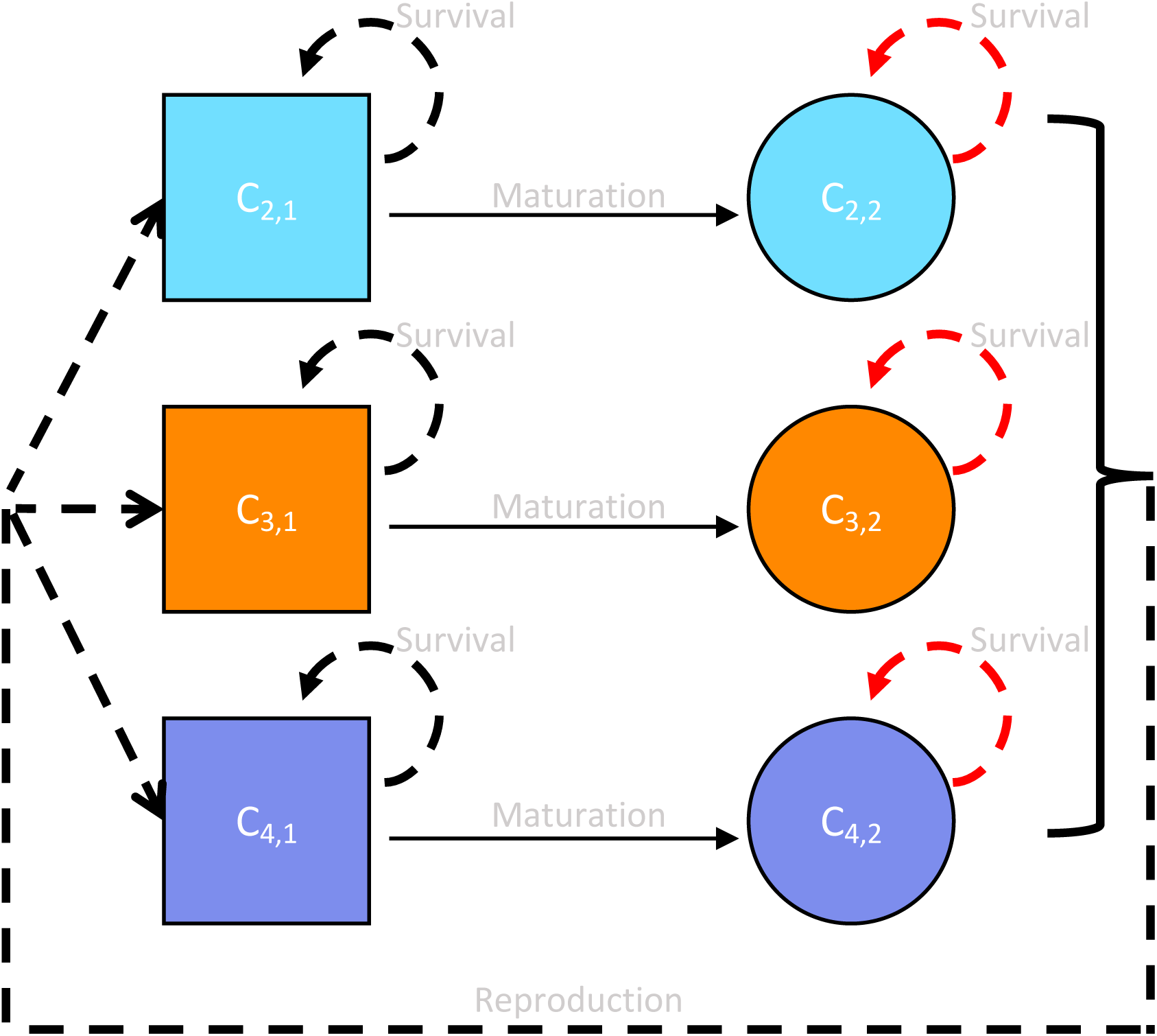
Simplified model conceptualization. This model includes two life-stages: immature (squares) and mature (circles), as well as three cytotypes: diploids (blue), triploids (orange), and tetraploids (purple). The number of individuals after each time step is determined based on three processes: reproduction, survival, and maturation. Arrows indicate relationships between the number of individuals of a stage at time *t*, and the number of individuals of a stage at time *t* + 1. Black dotted lines indicate density-dependent processes; red dotted lines indicate processes influenced by environmental stochasticity.

### Model description

The organism types in our model consist of three cytotypes (diploids, triploids, and tetraploids) at two life stages (immature and mature). The number of individuals in each of these six stages is denoted as *c_i, j_* (*t*) where the sub-index *i* = {2, 3, 4} keeps track of the ploidal level and *j* = {1, 2} the life stage (*i* = 1 indicates a reproductively immature individual, and *i* = 2 indicates a reproductively mature individual). The number of individuals at each stage at time *t* + 1, is determined by reproduction, survival, and maturation (depicted in a life-cycle graph in Fig. 1). We let *F_i,k_* (*t*) be the number of immature individuals of ploidal level *i* produced by mature individuals of ploidal level *k*. The number of individuals that survive to the next time step is determined by *S_i,j_*, the survival probability of individuals with ploidal level *i* and life stage *j*. Therefore, the number of immature individuals at the next time step of ploidal level *i* is *S_i,1_*(*t*) ∑_*k*_ *F_i,k_* (*t*), the total number of offspring of ploidal level *i* produced by reproductively mature individuals times their ploidy-specific survival probability. The number of reproductively mature individuals of ploidal level *i* at the next time step is simply those mature individuals that survive (with time-dependent probability *S*_2,_ *_j_* (*t*)) plus the immature individuals that mature (with constant probability *M_i_*). Accordingly, we can summarize these demographic transitions with the matrix

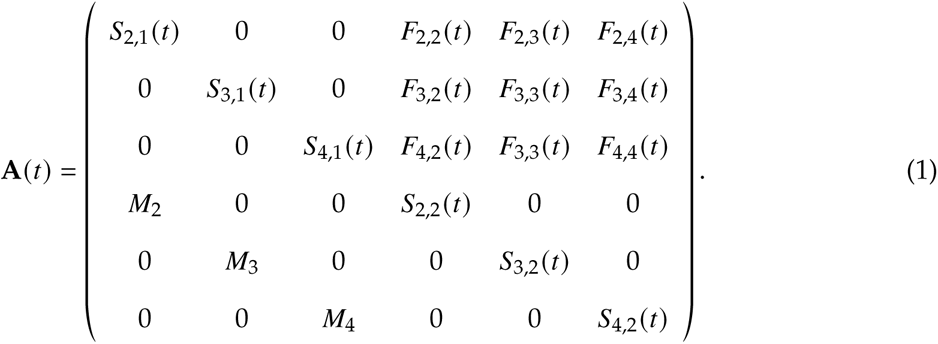

Using matrix notation, the dynamics of the number of cytotypes follows the recursion

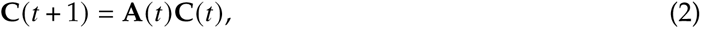

where **C**(*t*) = (*c*_2,1_(*t*), *c*_3,1_(*t*), *c*_4,1_(*t*), *c*_2,2_(*t*), *c*_3,2_(*t*), *c*_4,2_(*t*))*^T^*, the column vector of cytotype abundances.

The elements of the transition matrix **A**(*t*) are a mixture of time-independent constants and time-dependent functions of density and the environment. Here we detail density-dependent reproduction and survival at every stage by making the elements of the transition matrix functions of the densities of all types at each time step. Additionally, we let some of the functions be defined as random variables to incorporate demographic and environmental stochasticities. As such, the transition matrix **A** becomes a stochastic matrix. Although few analytical results are readily available for complex stochastic matrix population models, their formulation is amenable to exploring the statistical/probabilistic properties of the model solution, such as persistence probabilities of each type, which we define below in multiple ways. In what follows, we first describe how we formulated all the elements of this matrix having to do with reproduction and then all the elements having to do with survival and maturation.

#### Reproduction

We define the number of immature individuals of each cytotype produced by each mature individual as a stochastic process involving two main steps: gamete production followed by the union of those gametes (*i.e.*, reproduction). The details of the simulation process are visually and textually described in Figure S1. We assume that the total number of gametes produced by an individual of each cytotype may vary from year to year because of density-dependence and demographic stochasticities, but that during a single time step all individuals of the same ploidal level produce the same number of gametes. Demographic stochasticity and density dependence are entered in the total gametic production by drawing the total number of gametes *X_i_* (*t*) produced by an individual of type *i* out of a Poisson random variable. Specifically, we let

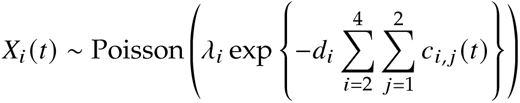

where *λ_i_* (*gnum* in our code) is the mean number of gametes expected by each individual in the absence of density dependence and *d_i_* is the strength of density dependence for each cytotype. As a note, in our figures, tables, and code, the vector of total gametic production ( *X*_2_, *X*_3_, *X*_4_) is denoted as *gam.vec*. The above description assumes that all members of a population, regardless of an individual’s cytotype or stage, will influence the number of gametes produced by each individual. Therefore, we define the number of gametes each mature individual produced by sampling Poisson distributions at each time step for diploids, triploids, and tetraploids independently across time and ploidal level.

We partitioned the total number of gametes into each potential type as follows. Of the gametes produced, unreduced gametes are produced by both diploids and tetraploids at frequency *b*. Surveys of naturally occurring populations have found the majority of populations surveyed have a low production of unreduced gametes (less than 2%); however, a minority of populations were found to only produce unreduced gametes (*i.e.*, 100%) (as reviewed by Kreiner et al. 2017*a*; Ramsey and Schemske 1998). Based on the natural population’s average rate, we set the frequency of unreduced gametes to 2% for our simulations, which we take as a moderate value. Due to the complexity of this model, we did not vary the frequency of unreduced gametes among individuals or between cytotypes, although such variation has been observed in naturally occurring autopolyploid populations (Kreiner et al. 2017*b*). Little is known about triploid gamete formation; however, only a small proportion of gametes produced by triploids are expected to be viable (Ramsey and Schemske 1998). Therefore, our model only allows triploid individuals to produce viable 3*n* gametes at the rate of *v*, set to 5.2% as defined by Ramsey and Schemske (1998).

After gamete formation, we allowed for a mixed mating system where both selfing and outcrossing can occur. The proportion of the total gamete pool that reproduce through selfing, is defined by *s*, which is historically known as the selfing rate. The remaining (*i.e.*, non-selfing) gametes outcross. It is well known that reproductive isolation between diploids and tetraploids could allow coexistence (Husband and Sabara 2003); therefore, as a proxy for reproductive isolation, we allow group-based assortative mating (Otto et al. 2008 and see Fig. S1) which we refer to as mating choice (mc). We define mating choice as the proportion of outcrossing gametes that unite only with gametes of the same type, and so mating choice takes values between 0 and 1. For example, when mating choice is equal to 0.1, then 10% of all diploid gametes that outcross will only unite with diploid gametes, and the remaining 90% will unite with gametes from any cytotype in the outcrossing pool.

In each assortative mating group, as well as in the outcrossing pool, we define the over-all probability of each offspring type based on the probability of sampling each gamete type. We then generate a random sample of offspring types (four possible types: diploid, triploid, tetraploid, or nonviable) using a multinomial distribution with sixteen categories (see Fig. S1). This approach allows for random union of gametes during outcrossing, which could allow for additional selfing to occur. Therefore, we calculate the probability of each offspring being formed from gametes derived from different parents. Then, we use a binomial distribution to sample the success of each offspring based on this probability to ensure the selfing rate is equal to the set rate, *s*. Finally, we only retain offspring with cytotype identities equal to diploid, triploid, and tetraploid (*i.e.*, 2*x*, 3*x*, 4*x*), assuming offspring of higher ploidal levels are nonviable (although this assumption may be violated in some systems).

#### Survival

While gametic production and reproduction incorporate only one stochastic process (demographic stochasticity), we chose to include both environmental and demographic stochasticities into the survival processes. Specifically, we assume that survival at the immature stage (*i.e.*, immature survival) is subject to demographic stochasticity, while survival at the mature stage (*i.e.*, mature survival) is subject to environmental and demographic stochasticities. Environmental stochasticity is expected to affect immature survival, and a future modeling effort could certainly incorporate it. Here, we added environmental stochasticity in only one of the life stages to be able to clearly tease apart its effects. A common observation and working assumption of models of iteroparous, perennial plants is that density-dependence acts most strongly at the seedling stage, with adults largely insensitive to density (Caswell 2000). Thus, we assumed that immature survival is density-dependent whereas mature survival is density-independent. We define the probability of survival for an immature individual of an individual of cytotype *i* as 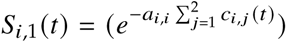, and assume survival of individuals of each cytotype is independent of any other cytotype in the population. Therefore, the number of immature survivors for each cytotype at a time step *t* follows a binomial distribution with “success” probability *S_i_*_,1_(*t*) and *c_i_*_,1_(*t*) trials. Immature survival is density-dependent because the survival probability is itself a declining function of density.

Mature survival was defined as a binomial trial with an environmentally determined survival probability, which we assumed was independent and identically distributed (*iid*) over time. This specification makes mature survival a stochastic process with both demographic and environmental stochasticities. This binomial distribution with *c_i_*_,2_(*t*) trials and time-dependent survival probability, *S_i_*_,2_(*t*), which is itself a random variable. The survival probability varies over time following a Beta distribution, which is a flexible distribution that takes values between 0 and 1. We sampled survival probability independently over time and across cytotypes, which implies that each cytotype perceives the same environmental state differently. We parameterize the Beta distribution in terms of its mean, *µ*, and variance, *cr*^2^ (see Kortessis and Chesson 2019 for details). When written this way, the traditional shape (*α*) and scale (*β*) parameters are 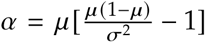 and 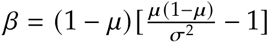 such that the mean and variance can be manipulated independently. However, because of the bounded range of the Beta distribution, the variance can be changed independently of the mean only within some bounds. Specifically, *cr*^2^ *< µ*(1 − *µ*) (Morris and Doak 2004), meaning that a limitation exists on the amount of environmental variance that can be explored for different mean survival probabilities, *µ* (Table 1). To get around this constraint, we considered different levels of environmental variability by defining *ci* = *cr*^2^/[*µ*(1 − *µ*)] (0 ≤ *ci <* 1), the proportion of the maximum possible environmental variance. The parameter *ci* can still be interpreted as the level of environmental variability because *ci* is proportional to the variance; when *ci* is small, the variance in mature survival probability is small, regardless of mean survival probability.

**Table 1:**
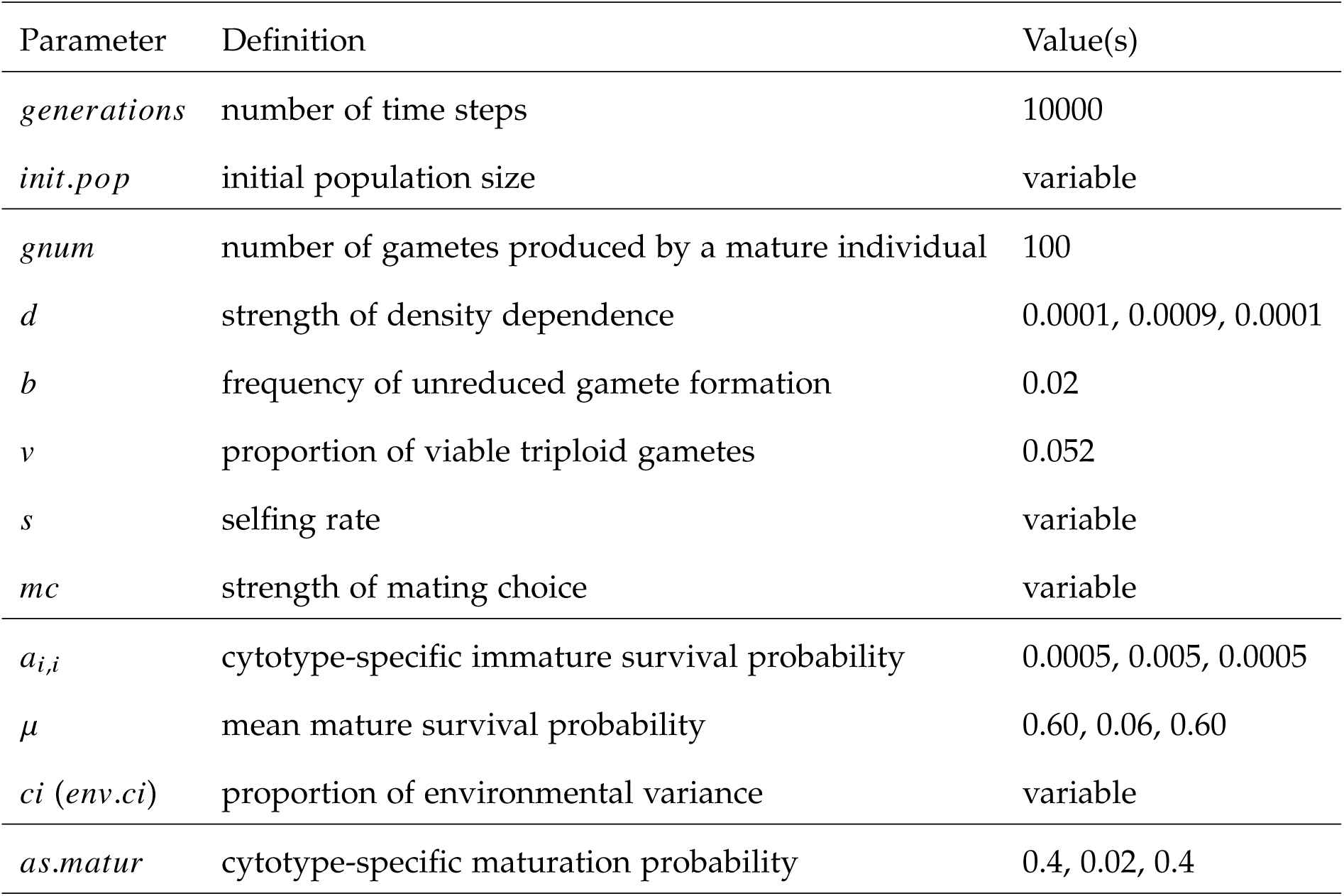
Summary of model parameters, their definitions, and values. If three values are indicated, the value is cytotype-specific and ordered as diploid, triploid, and tetraploid. If the value is indicated as variable, we investigated multiple values for this parameter as described in the Simulation Settings section of the Methods.

#### Maturation

The number of immature individuals that mature at each time step was sampled independently for each cytotype using a binomial distribution where the number of trials was equal to the number of cytotype-specific immature individuals at time *t*, and the probability was set as a cytotype-specific maturation probability (denoted as *as*.*matur* in our Figures, tables and code).

### Simulation settings

We investigated 1089 different parameter combinations to characterize the effects of reproductive isolation, founding population size, and variation in the environment on coexistence and stability among mixed cytotypes. These 1089 parameterizations consist of every possible combination of selfing rate (0 - 1, by 0.1; 11 values), mating choice (0 - 1, by 0.1; 11 values), initial population size (10, 100, 1000; 3 values), and proportion of environmental variance (0.1, 0.5, 0.9; 3 values) (Table 1). Each parameter set was used to simulate the overall population dynamics for 10000 generations and was repeated for 500 replicates, yielding 1089 × 500 = 544, 500 replicate simulations. Our initial population size, mimicking a colonization founding event, consists only of mature diploid individuals. All the remaining model parameters were left unchanged across the 1089 parameterizations (Table 1). Based on our focal questions, we assumed diploids and tetraploids had equal mature and immature survival rate, maturation rate, and strength of density dependence. As such, the type with greater reproductive ability is a dominating competitor. Triploids, on the other hand, were set to have lower mature and immature survival rates than diploids and tetraploids, as expected in nature (Felber and Bever 1997; Husband 2004). Maturation rate was also assumed to be lower in triploids compared to diploids and tetraploids. For triploids, the strength of density dependence was set as greater than for diploids and tetraploids.

### Simulation Output

#### Population Composition and Coexistence

We sought to characterize the long-run trajectory of diploids and tetraploids for every parameter combination and model replicate via different summary statistics. These summary statistics were computed for all the combinations of selfing rates, mating choice, initial population size, and strength of environmental stochasticity.

To compare general population trajectories among the different parameterizations, we visually inspected the joint abundances of diploids and tetraploids after an initial “burn-in” period of 20 generations to identify if there are general trends or apparent basins of attraction for the model solutions.

We assessed the variation in the resulting frequency of simulations where we observed coexistence, extinction, and fixation (defined below) among the different parameter combinations and model replicates. A general assumption of similar models typically used to study population growth and selection is that one tracks abundance with population density (*e.g.*, abundance per unit area), which lets population sizes change continuously (*e.g.*, Lotka-Volterra competition models). Extinction in such models of competition similar to ours is then defined as a long-run trajectory of population density towards zero (despite never actually reaching zero). Moreover, these studies typically consider only two types, in which case extinction and fixation are two sides of the same coin: when one type goes extinct, the other “fixes”. In population models such as the one used here, it is possible for both cytotypes to go extinct. Thus, extinction and fixation manifest differently. Those differences help us tease apart the environmental drivers that explicitly contribute to extinction. For all 500 replicates for each model type, we calculated the mean abundance of all diploids and all tetraploids (*i.e.*, both immature and mature) for 500 time steps after burn-in. We then calculated the geometric mean of those means as the square-root of the diploids’ temporal mean times the tetraploids’ temporal mean for each replicate. We define coexistence at each replicate as both the diploid and the tetraploid being present at a mean relative abundance equal to or greater than 0.1. Therefore, we defined a coexistence cutoff equal to 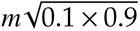, where *m* is the sum of the mean of diploids and tetraploids. Because these calculations can potentially hide times of low abundance, we also calculated an alternative measure by replacing the mean abundance of each cytotype with their geometric means over time, which is more influenced by small values. Based on the first measure, we identified the probability of extinction and fixation as a function of mean abundance of each cytotype. Extinction was defined as a cytotype-relative mean abundance below or equal to 0.1, while fixation indicated if the relative mean abundance was above or equal to 0.9. We use cutoff values for extinction and fixation because the extinction states (zero abundances) are not absorbing in our model. Biologically speaking, a cytotype may reach zero abundance but re-emerge later due to formation of gametes produced by other cytotypes that are still present in the population. These nuances are particular to systems like ours, which although general, seldom have found a formal treatment in population dynamics theory. Hence, we examined extinction, fixation, and coexistence and their relationship to cytotype mean abundance.

#### Stability

We define stability as the entire population’s response to environmental noise. In the face of the same amount of variation in the quality of the environment, a stable population will not experience large fluctuations in population size whereas an unstable population will. As extensively shown and studied by Ives et al. (2003), the processes that determine the response of the population to environmental variation are intra- and inter-specific competitive effects (*i.e.*, density-dependence).

Stochastic population dynamics models conceptualize environmental fluctuations as exogenous forces affecting the growth rate of a given taxon. The quality of the environment is directly translated into a positive or a negative effect on the growth rate of this species. Such translation involves intrinsic properties of the dynamics of the taxon, such as its maximum growth rate and the strength of intra- and inter-specific density-dependence. In a very real and practical sense, the temporal fluctuations of population abundances can be seen as the outcome of an external input (the environment) being fed into a system (the population). Stochastic population dynamics theory and practice have shown us that it is those intrinsic properties that modulate the translation of the environmental fluctuations into growth rate fluctuations (Ives et al. 2003). Weaker density-dependence generates stronger reactions to environmental variability. This strong reaction manifests itself in the form of greater variability of the total population size. Strong density-dependence generates weak reactions to environmental noise. In the face of the same environmental variability, a population with a strong density-dependence will fluctuate less than a population with a weak density-dependence. For a single taxon, stability can be measured as the ratio of the magnitude of environmental variation to the strength of density-dependence, which allows for a direct comparison of the reaction of two different taxa to the same environmental noise regime.

Ives et al. (2003) findings imply that deeming a particular set of time series as representative of stable or unstable dynamics just by its overall variability and aspect might be misleading and conflate the fundamental processes governing the dynamics of an ensemble of interacting populations. Therefore, we used the theoretical framework of Ives et al. (2003) to estimate four different stability metrics that in essence, and without entering into mathematical details (see help files of our R code), give a standardized measure of the response of a community to environmental noise. The stability metric of choice, the variance proportion at stationarity due to environmental variation, or simply variance proportion, measures how the long-run variance of the total population (*i.e.*, composed of all the cytotypes) compares to the variance of the environmental noise process (Ives et al. 2003). In a highly unstable system, the environmental noise variability is translated into a greatly amplified variance of the long-run variance of the population and this metric will be close to 1. In a highly stable system, the environmental noise variability is not translated into amplification, and thus the variance proportion will be equal to 0. To obtain these stability metrics, we first fitted the model of Ives et al. (2003) to the the multi-types (cytotypes) time series simulations for all the model parameter combinations using their conditional least squares approach (Ives et al. 2003). This model corresponds to a multi-species or multi-types stochastic population dynamics model with intra- and inter-specific density-dependence and environmental variability; specifically, a multivariate stochastic Gompertz population dynamics model under environmental variablity. Despite our simulation model being decidedly more complex than the Gompertz model, evidence from prior studies shows that the simple stochastic Gompertz model is nonetheless a powerful tool to summarize elementary properties of the dynamics of natural populations, and that these inferences are robust to departures from the model assumption (Ponciano et al., 2018). Based on an abundance matrix for diploids and tetraploids for 500 time steps following burn-in, we estimated how the abundance of one cytotype influences the growth rate of the other cytotype.

For all model replicates, a growth rate covariance matrix of all generations after burn-in of diploids and tetraploids was calculated. This covariance matrix represents the extent to which the growth rates in diploids and tetraploids change together, as well as the variance in growth rates for diploids and tetraploids. We examined the distribution of model replicates for the first value of the eigenvector of this covariance matrix. The first value of the eigenvector represents the main principal component of the data; the larger this value, the more variance in the data set. In addition, we looked at the determinant of the covariance matrix, which represents the generalized variance.

Finally, we translated this simulation framework into R software that is available via our package AutoPop (https://github.com/mgaynor1/AutoPop). Scripts associated with model evaluation and generation can be found in DRYAD

## Results

The first notable result is that changing initial population size did not influence the statistical properties of the population trajectories. Although future analyses may include a larger range of the initial population size, a number of clear conclusions emerge from the numerical experiments.

### Population Composition and Coexistence

At low environmental variance, populations consistently converge to what seemingly appear as two distinct attractors (Fig. S2, light blue dense areas), regardless of parameters related to reproduction: (1) high abundance of diploids and low abundance of tetraploids; or (2) low abundance of diploids and high abundance of tetraploids. As environmental variance increases, both cytotypes are more likely to become extinct, and the distinction between these attractors vanishes. This distinction also diminishes as selfing rate or mating choice increases (Fig. S2 and Fig. S3, respectively).

Extinction, fixation, and coexistence probabilities all changed as a function of the mean abundances at stationarity and selfing rate (Fig 2). Extinction was unlikely for both diploids and tetraploids at high selfing rates and high values of mating choice (Fig. 2A and 2D, Fig. S4A and S4D, dark red points). However, when selfing is rare, tetraploid extinction probability was high whereas diploid extinction probability remained low (Fig. 2A and 2D; black or dark blue points). Furthermore, under low selfing rates mean abundance of tetraploids was low, whereas mean abundance of diploids was high. Such divergent shifts in average abundance are reflected in the fact that fixation rates were high for diploids and low for tetraploids at low selfing rates, and that extinction and fixation rates were uniformly low at high selfing rates. Additionally, coexistence is only likely under high selfing rates, when the probability of neither extinction nor fixation is very high (Fig 2C and 2F).

**Figure 2:**
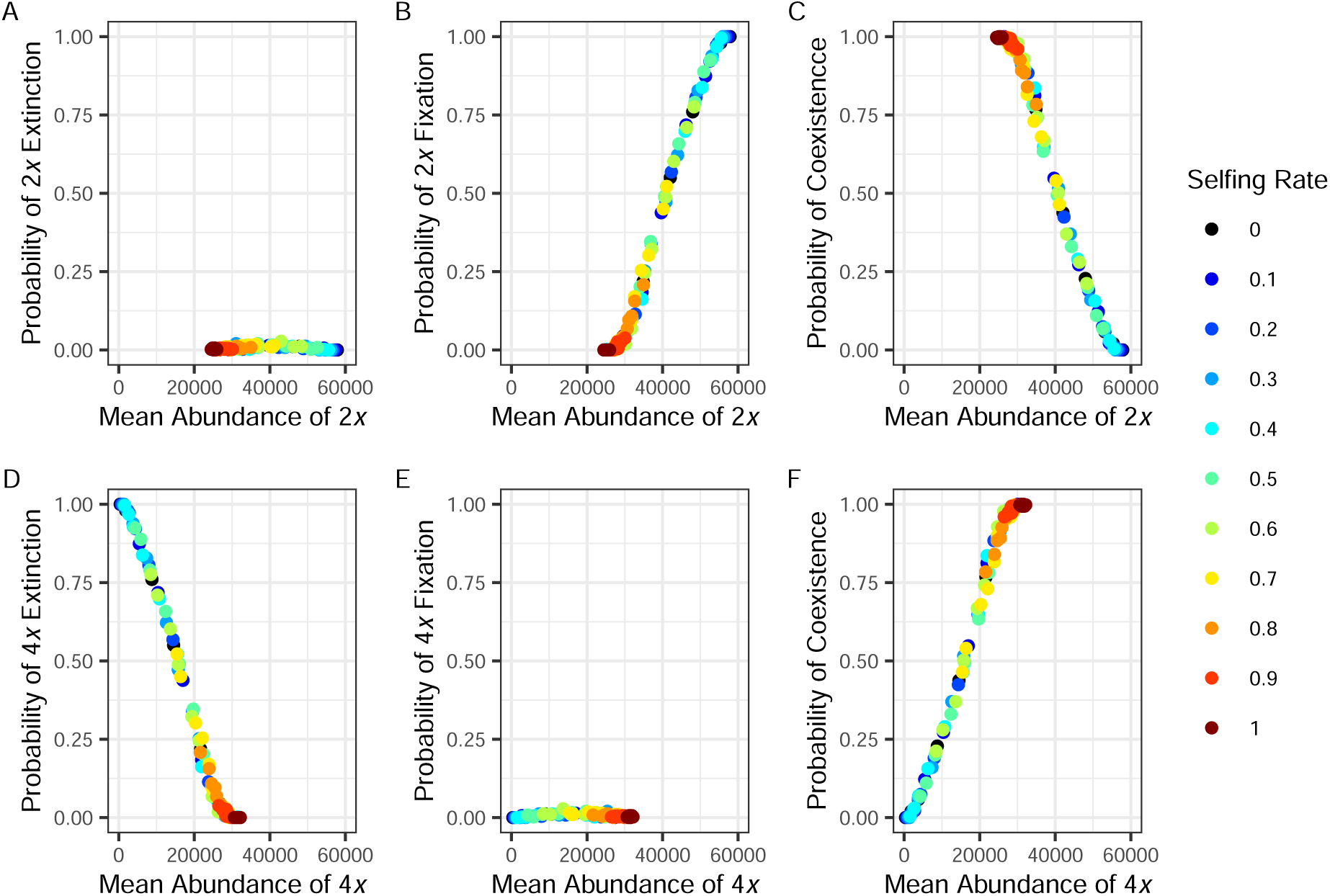
Probability of extinction (A, D), fixation (B, E), and coexistence (C, F) as a function of the mean abundance of each cytotype for 500 generations after reaching stationarity. For tetraploids, increased selfing rate is observed with increased mean abundance and decreased probability of extinction. For diploids, increased selfing rate is related to decreased mean abundance after stationarity; lower selfing rates increase the probability of fixation.

Decreased mating choice typically reduces the probability of coexistence (proportion of the simulations under a given setting where coexistence occurs), although this effect is diminished by selfing rate and environmental variability. At low environmental variance with no selfing (Fig 3, upper leftmost panel), increasing the mating choice increases the likelihood that diploids and tetraploids coexist. Under moderate selfing (0 *< s <* 0.5), effects of mating choice on coexistence of cytotypes are muted (Fig 3, center column of the top row), and no discernible effect of mating choice under complete selfing (*s* = 1) (Fig 3, rightmost column of upper row). When environmental variance increases (Fig 3, middle row), we observe higher variance over time in the mature survival probability which allows for both a higher proportion of coexistence outcomes (a higher density of the saturated colored histograms) and a higher variability of the equilibrium geometric mean of the two population sizes. Roughly, an increase in the temporal variation of the environment, reflected in an increased variation of the survival of mature individuals, equalizes the chances of coexistence for all mating choice values as well as for all selfing rate values. Remarkably, although the long-run geometric mean of the two types seems to be uniform under intermediate environmental variability, the geometric mean of the two types in the instances where coexistence does not occur (the transparent section of the histograms) becomes more variable. Finally, a very high increase in environmental variance allows for more population crashes and hence dramatically increases the proportion of simulation results without coexistence (Fig 3, third row). With high environmental variance, coexistence occurs at a much lower geometric mean than at lower environmental variance. These observations were consistent for our alternative calculations that account for drastic abundance shifts (Fig. S5).

**Figure 3:**
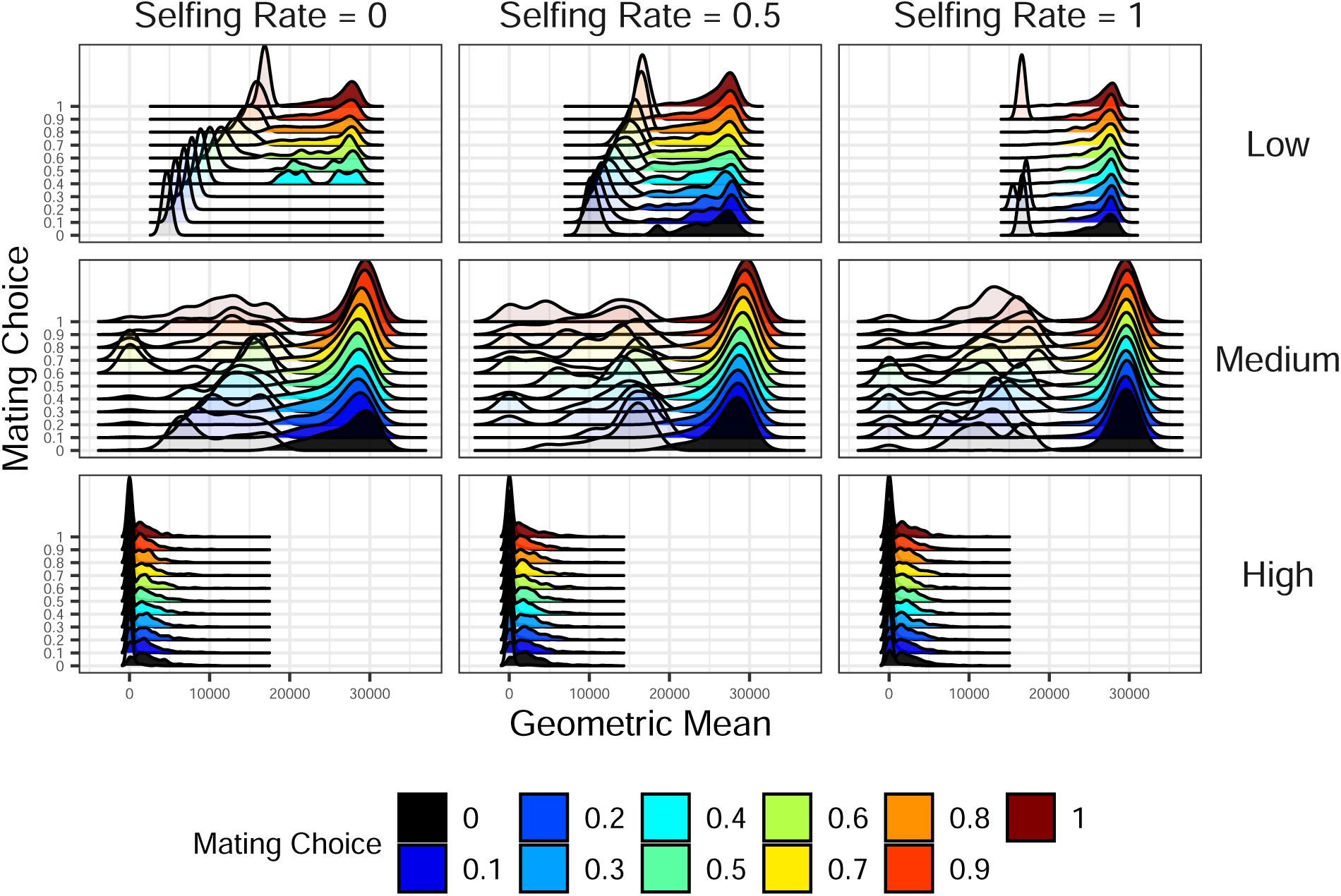
Smoothed histograms of the geometric mean of the population trajectories for both diploid and tetraploid population sizes for 500 generations after burn-in across all models with an initial population size of 100. Because coexistence is defined as diploids and tetraploids each existing above a threshold of a 10% proportion, this threshold corresponds to different geometric means under the different simulation settings. However, a common cut-off coexistence threshold above which the cytotypes coexist and below which they fail to coexist can be computed (see text for details). This calculation allows splitting each smoothed histogram of the final geometric mean into two sides: geometric means below the threshold (transparent coloring on the left) and those above the threshold (fully saturated color on the right). As environmental variance increases, variability in coexistence increases. At high environmental variance, coexistence occurs at a lower geometric mean, suggesting coexistence is less secure.

### Stability

Using our simulation framework to study the dynamics of a population composed of multiple cytotypes allows us to understand how the properties of this system interact with extrinsic environmental variability. The first conclusion from our simulations is that reproductive isolation, more than selfing rate alone, is key to generating stability of the long-run population size of diploids and tetraploids. Our simulations show that as the environmental variance increases, selfing rate ceases to be a determinant of the stability of the long-run population sizes (Fig 4). At low environmental variance, however, higher selfing rates and increased mating choice increases the stability of the long-run population size. Effectively, as selfing rate or mating choice goes to 1 (*i.e.*, reproductive isolation increases), the stability of the system increases.

**Figure 4:**
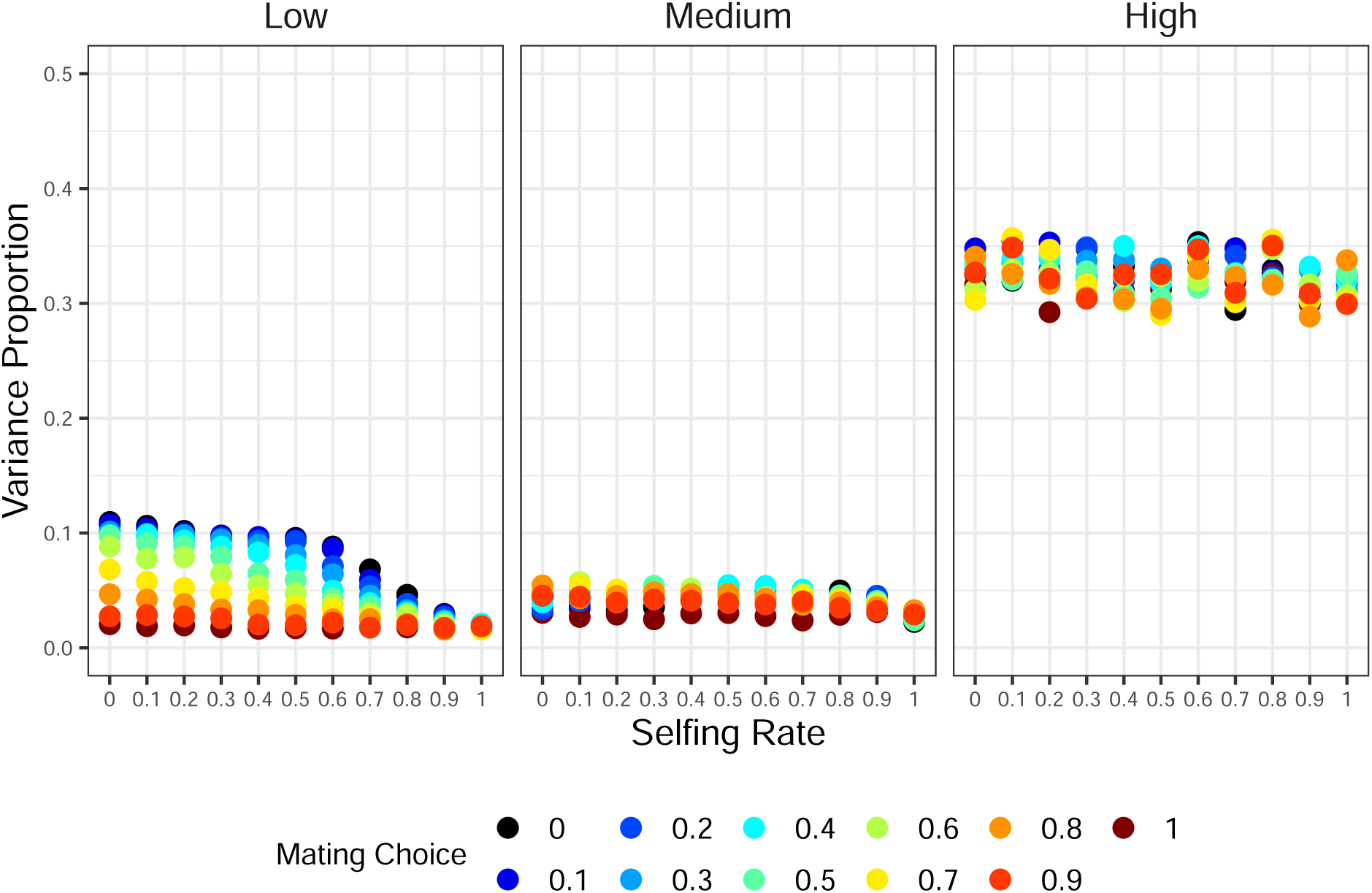
Changes in a stochastic stability metric as a function of mating choice and selfing rate for all models where initial population size is equal to 100. The variance proportion attributable to environmental noise at stationarity is a standard stability metric in stochastic systems (Ives et al 2003). The lower values of the variance proportion, the more stable the system. Here, increasing the mating choice and the selfing rate is expected to result in an increased stability when environmental variance is low or medium.

The second conclusion, drawn from a visual inspection of diploid and tetraploid intraspecific density-dependence, is that as environmental variance increases, the strength of intraspecific competition (*i.e.*, self regulation) diminishes (Fig. S6, Fig. S7). This tendency is much stronger in diploids than in tetraploids. For diploids under low environmental variability, a low mating choice (or selfing rate) is translated into strong intraspecific competition effects, whereas a high mating choice results mostly in strong to weak density-dependence. As environmental variability increases, the difference between mating choice values gets diluted, and the strength of the self-regulation of the diploid population wanes (as seen as all the densities piled up between 0.5 and 1 on the center panel and upper row of Fig. S6). Because environmental variability affects the maximum growth rate of the population (which is roughly births minus deaths) via the death rate, as the environmental variance becomes higher, the relative importance of the maximum growth rates becomes higher *vis-á-vis* the density-dependent factors.

Lastly, as environmental variance increases, the outcome of competition between diploids and tetraploids, measured by their differential growth rates, shifts to favor tetraploids (Fig 5). In the near absence of environmental variability, the deterministic processes, such as the strength of intraspecific regulation, and stochastic demography properties fully determine the outcome of the competition between the cytotypes. In short, a nearly constant environment allows for cytotype-specific growth rates, especially when mating choice and selfing rates are low. When both cytotype reproductive isolation (*i.e.*, mating choice) and environmental variance are low, and selfing rate is either null or 0.5, diploids have clearly higher growth rates (Fig 5, black and dark blue points on the upper left panel). However, as cytotypes become more reproductively isolated, we observe an agglomeration of simulation results around the following outcomes: both cytotypes (diploids and tetraploids) have approximately equal growth rates, diploids have a higher growth rate than tetraploids, or tetraploids have a higher growth rate than diploids. Finally, when environmental variance is high, these trends are almost completely undone. As environmental variance increases, the growth rate of the tetraploids is always higher than that of the diploids; however, the extent of this advantage varies (Fig 5). When cytotypes are reproductively mixing (selfing = 0 and mating choice is low), it appears that diploids are dominant, but this fact becomes highly uncertain across replicate populations in more variable environments. Thus, it appears that increasing reproductive isolation between cytotypes (*i. e.*, increasing selfing and mating choice) and increasing environmental variation tends to reduce inequalities in intrinsic growth rates.

**Figure 5:**
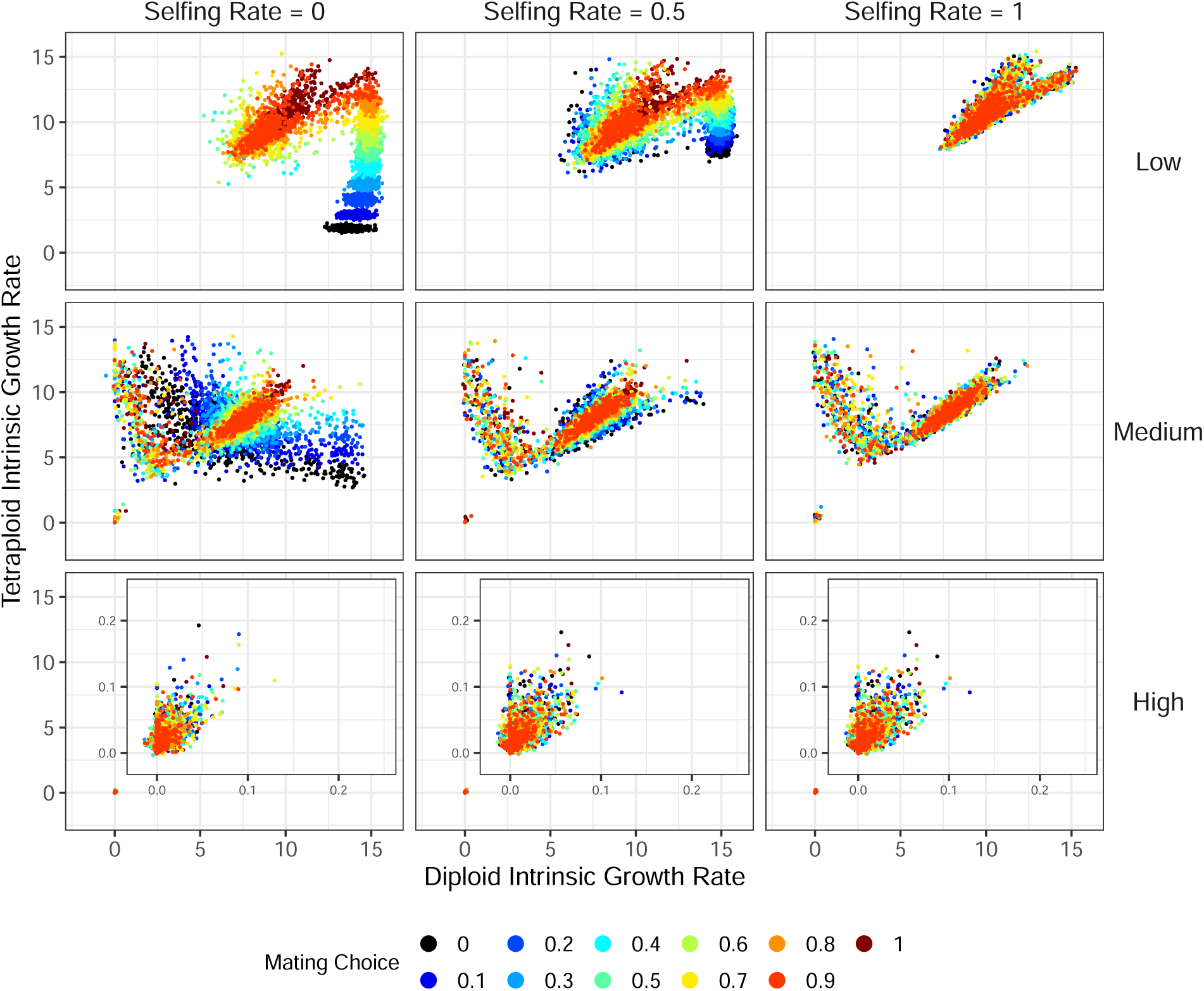
Intrinsic growth rate, as derived based on conditional least squares, for low, medium, and high environmental variance where the initial population size is equal to 100. Each point represents a model replicate; the agglomeration of points indicates variance in cytotype intrinsic growth rate by chance. Overall, diploid and tetraploid growth rate decrease with increased environmental variance. In low environmental variance we observe neutral competition, diploid competitive advantage, and tetraploid competitive advantage. In medium and high environmental variance, we observe only a tetraploid competitive advantage; however, the strength of this advantage varies.

## Discussion

In this study, we seamlessly connect stochastic population dynamic principles and empirical understanding of mixed-cytotype populations to improve our understanding of the formation, establishment, and persistence of tetraploids. We developed a novel stochastic matrix population dynamic model for long-lived diploids, triploids, and tetraploids with ongoing gene flow. Through this model, we elucidate the conditions under which coexistence and stability of mixed-cytotype populations are possible. This model includes, by design, known factors promoting coexistence, such as recurrent formation, niche independence, and stochasticity. Based on the response of our simulated mixed-cytotype populations to environmental fluctuations, we found stability is supported by reproductive isolation through increased selfing and increased assortative mating. We also found that the observed frequency of each cytotype in a population may not be indicative of dynamics among cytotypes, because chance alone determines the dominant cytotype at low environmental variance. Additionally, we found that at higher environmental variance, tetraploids have a larger competitive ability compared to diploids.

### Coexistence

Under recurrent stochastic formations, as well as equal and independent survival probabilities, this model favors coexistence of mixed cytotypes. This model also focuses on an iteroparous perennial system with overlapping generations, as perenniality is common in polyploids (Stebbins 1950; Thompson and Lumaret 1992; Van Drunen and Husband 2019); by definition, more instances of reproduction associated with a perennial life history will increase the probability of successful offspring, and therefore escape from MCE. Perennial life history has been theoretically shown to increase the probability of polyploid establishment (Van Drunen and Friedman 2022).

In our model, we allow consistent unreduced gamete formation over time, thus continued formation of autotetraploids and increased probability of persistence by chance (Soltis and Soltis 1999). Recurrent formation has been inferred for numerous extant naturally occurring autotetraploids (Husband and Sabara 2003; Levin 2011; Parisod and Besnard 2007; Ramsey et al. 2008; Segraves and Thompson 1999; Servick et al. 2015; Těšitelová et al. 2013), and previous theoretical models have found that recurrent formation supports persistence of the autotetraploid (Baack 2005; Felber 1991; Oswald and Nuismer 2011; Rausch and Morgan 2005; Rodriguez 1996). Due to the complexity of our model, only a single rate of unreduced gamete formation was allowed among individuals and across generations. However, unreduced gamete formation in naturally occurring populations is poorly investigated, and the extent to which unreduced gamete production varies temporally is unknown. Thus, incorporating temporal dynamics of unreduced gamete production into our model, while interesting, would involve the use of arbitrary values. Because production of unreduced gametes has been suggested to increase due to stress and a fluctuating environment (Bomblies and Madlung 2014; Kreiner et al. 2017*a*; Ramsey and Schemske 1998), temporal variance in unreduced gamete production could be achieved in future studies through incorporating environmental stochasticity into the probability of unreduced gamete formation.

Our model includes niche independence due to the mature survival probability of each cytotype having an equal, but independent, relationship to environmental stochasticity; although cytotypes maintain an equal probability of survival, they experience different patterns of survival over time and perceive the same environmental state differently. By allowing survival probability to be independent among cytotypes, divergence in niche is possible due to a storage effect (Chesson and Warner 1981; Kortessis and Chesson 2021). Niche divergence has been considered essential for an autotetraploid to overcome MCE and establish (Levin 1975, 2003). Despite the expectation of niche divergence, lack of divergence in broad-scale climatic niches has been observed in multiple mixed-cytotype autopolyploid systems (Chung et al. 2015; Diaz 2020; Gaynor et al. 2018*a*; Godsoe et al. 2013). Additionally, past theoretical models have included explicit niche divergence through competition with their diploid progenitor (Fowler and Levin 1984; Rodriguez 1996); however, niche differentiation could be driven by additional interactions and community dynamics (Gaynor et al. 2018*b*; Segraves and Thompson 1999). For example, broad-scale niche conservatism was observed among diploid and autotetraploid *Heuchera cylindrica* (Saxifragaceae) (Godsoe et al., 2013), but further analysis identified potential differentiation in nutrient acquisition due to divergent interactions with below-ground mutualists (Anneberg and Segraves, 2019). Recent work suggests that, in some instances, cytotypes can perceive the same environmental state differently (Bafort et al. 2023). Hence, instead of modeling explicit directional competition among cytotypes and their community members, our study allows divergence in niche among cytotypes to be driven by the environment. To our knowledge, this is the first study to allow environmentally driven niche divergence between cytotypes, which is commonly hypothesized for maintaining species diversity in plant communities (Angert et al. 2009; Usinowicz et al. 2017).

Incorporating stochasticity in our model creates opportunity for cytotype persistence through ’stochastic flickering’. In deterministic models, we expect a single equilibrium to be reached, as seen in many previous theoretical investigates of MCE (Burton and Husband 2001; Felber 1991; Felber and Bever 1997; Fowler and Levin 1984; Levin 1975). In stochastic models, however, deterministic equilibria no longer exist. A single stable equilibrium under stochasticity is translated into a stationary, or quasi-stationary, distribution around which a population fluctuates, which can be described as a stochastic attractor (Allen 2010.) When converted into a model with environmental stochasticity, a deterministic model that exhibits multiple equilibria may exhibit a behavior, by chance, in which the population trajectories may jump between the different attractors. This random movement between attractors under environmental stochasticity is termed ’stochastic flickering’ (Gardner et al. 2000; Ponciano and Capistrán 2011; Touboul et al. 2018). For each cytotype, the movement between equilibria is not synchronized; therefore, there are stretches of time where one cytotype dominates the other and vice versa. The inconsistent timing in flickering therefore provides opportunity for the minority cytotype to gain advantage over the other and become the majority cytotype. Therefore, stochasticity can lead to maintenance of multiple cytotypes.

### Stability

#### Reproduction

Increased selfing and increased mating choice both tend to stabilize population dynamics and promote coexistence of diploids and tetraploids in our model. Here we describe the caveats to our reproduction function. Additional dynamics known to influence cytotype persistence were not incorporated in our modeling framework due to the existing complexity of reproduction. Our model incorporates mixed-ploidy-specific gamete formation and gamete union based on the following five user-defined factors: the mean number of gametes per individual, a rate of unreduced gamete formation, the proportion of viable triploid gametes produced, selfing rate, and mating choice. These factors are sufficient to incorporate pre-zygotic barriers without having to define a large number of additional parameters. Also, group-based associative mating, as defined by mating choice, allows the rate of intra-cytotype gamete exchange to be defined without explicitly defining pollinator transitions, spatial distance, or other demographic factors. Previous spatially explicit theoretical work suggests the probability of establishment of a polyploid is influenced by three factors; local majority status (Baack 2005; Griswold 2021; Kauai et al. 2023; Spoelhof et al. 2020), pollen and seed dispersal probability distribution shape (Van Drunen and Friedman 2022), and clonal ability (Levin 2021; Spoelhof et al. 2020; Van Drunen and Friedman 2022). Of these, our model can incorporate the first two *via* our mating choice parameter. The impacts of clonality remain untreated by our modeling efforts.

Our model parameters define which gametes may unite, rather than the success of each offspring (*i.e.*, pre-zygotic rather than post-zygotic barriers). In nature, post-zygotic barriers are known to decrease the number of intra-cytotype offspring produced (Ramsey and Schemske 1998; Schneider et al. 2023; Sutherland and Galloway 2017; Sutherland et al. 2020). Post-zygotic barriers among cytotypes include genomic imbalance and meiotic irregularities (Kö hler et al. 2021, 2010; Sutherland and Galloway 2017); however, the strength of these barriers varies, and the barriers are often incomplete (Sutherland and Galloway 2017; Sutherland et al. 2020). Therefore, post-zygotic barriers could be included through defining viability of intra- and inter-cytotype offspring in future models.

When interpreting our findings, it is important to consider that we assumed selfing rate and unreduced gamete formation to be equal for diploids and tetraploids. Although selfing rate has been observed to be equivalent between diploids and autotetraploids with mixed mating systems (Ozimec and Husband 2011), shifts toward increased selfing in polyploids is both predicted (*e.g.*, Hedrick 1987) and observed, often involving loss of self-incompatibility (Cook and Soltis 1999; Husband et al. 2008; Sutherland et al. 2018). Under equal selfing rates between diploids and tetraploids, our model yielded opposite relationships between selfing rate and persistence for diploids and tetraploids. Specifically, our simulations show that an increased selfing rate decreases the probability of autotetraploid extinction, allowing an autotetraploid to escape MCE. In contrast, reduced selfing rates increased the probability of diploid fixation. This increase is likely only due to the low abundance of autotetraploids, because at this state diploids are expected to experience strong intraspecific density-dependence. Although selfing is known to aid newly synthesized polyploids, continued selfing may lead to inbreeding depression in both diploids and autotetraploids. Previous theoretical models have suggested that for autopolyploid persistence, inbreeding depression needs to be low or coupled with increased autotetraploid survival probability (Rausch and Morgan 2005; Van Drunen and Friedman 2022). However, autotetraploids will likely experience lower inbreeding depression compared to their diploid progenitors (reviewed by Clo and Kolář 2022), and inbreeding depression therefore will likely not impede establishment or persistence (Layman and Busch 2018). In addition to potential inbreeding depression, a recent theoretical study concluded that selfing may be less beneficial to autotetraploid establishment when polyploidy is associated with a fitness cost. This reduced benefit is due to self-fertilization selecting against genetically linked unreduced gamete formation (Clo et al. 2022). Similar to our model, these authors specified equal selfing rates for diploids and autotetraploids; therefore, it is unknown if this outcome is a result of these rates being linked.

#### Population Trajectory

The frequency of each cytotype observed in a population at one point in time only captures an instant of the inter- and intra-cytotype dynamics, yet the intricate processes resulting in one or the other type dominating or coexistence ultimately unfold over many generations. Temporal monitoring of perennial mixed-cytotype populations is rare, with the single set of long-term population surveys revealing stable cytotype frequencies over a 40-year timespan (Trávńıček et al. 2010). The paucity of systems with temporal surveys is likely due to the tedious nature of quantifying ploidal level and known extended life-span of multiple cytotype systems (Kolář et al. 2017; Van Drunen and Friedman 2022). Due to the stochasticity expected in naturally occurring populations, it is important to consider the history of the population rather than a single snapshot. For instance, chance may in some cases be the sole determinant of the dominant cytotype at low environmental variance. Specifically, simulation replicates experienced substantial variability in the competitive abilities of diploids and tetraploids, as defined by the intrinsic growth rate. Therefore, to extend the utility of this model to understand the history of extant mixed-cytotype populations, in an ongoing project, we are currently incorporating genotype.

#### Environmental Stress

Autotetraploids are more likely to persist than their diploid progenitors under high environmental variance. Survival of tetraploids under fluctuating environments is not unexpected, as polyploids are proposed to have persisted at environmental extremes (Brochmann et al. 2004; Ehrendorfer 1980), as well as through the Cretaceous-Paleogene (K-Pg) boundary (Fawcett et al. 2009; Fawcett and Van de Peer 2010; Levin and Soltis 2018; Vanneste et al. 2014). The K-Pg persistence has historically been attributed to various individual-based dynamics, such as genetic, physiological, and developmental change (reviewed by Levin and Soltis 2018; Van de Peer et al. 2021). Our simulations revealed that under higher environmental variance, tetraploids will have a larger competitive ability compared to diploids, despite equal survival probabilities. Here, we speculate that tetraploid persistence in fluctuating environmental conditions can likely be attributed to gamete “stealing”, shifts in strengths of density dependence, and niche differentiation.

The higher net growth rate observed for tetraploids compared to diploids is due to tetraploids experiencing a relatively weaker net density-dependent effect than diploids. The reproductive step in our model allows for gamete “stealing” (*i.e.*, diploid gametes producing tetraploid offspring) through recurrent formation and a triploid bridge. The net effect of this process is to increase the gains in the growth rate of tetraploids, hence diminishing the relative effect of density-dependence. Persistence of tetraploids through harsh environmental conditions has been suggested to be aided by their selfing ability (Freeling 2017), similar to our findings at medium environmental variance. However, under high environmental variance, tetraploid persistence is strongly determined by randomness, so much so that the outcomes cease to be driven by parameter values for selfing rate and mating choice. The tetraploid competitive advantage retained during high environmental variance is likely driven by continued primary contact through gamete “stealing”. As the majority of theoretical investigations into mixed-cytotype persistence and establishment have not included triploids and ongoing gene flow, these dynamics have been overlooked.

Previous theoretical investigations suggested that gamete “stealing” would not be sufficient to overcome MCE in conjunction with stochastic demographic and environmental processes (Fowler and Levin 1984; Levin and Soltis 2018); to overcome stochasticity, tetraploids would need to shift in growth investment or increase propogaule pressure (Levin 2021). However, growth rates in mixed-cytotype populations have rarely been compared. Similar to our theoretical result, the strength of density-dependence, in relation to neighborhood crowding, was found to be weaker for tetraploids than diploids in *Artemisia tridentata* (Asteraceae) (Zaiats et al. 2021). When crowding was absent, diploid growth rate was higher than the tetraploid growth rate; on the other hand, under crowding, this dynamic shifted, and tetraploids retained a competitive advantage (Zaiats et al. 2021). Similarly, in low environmental variance, we identified shifts in competitive ability among model replicates. In relation to environmental variation, the growth rate and carrying capacity of tetraploid *Spirodela polyrhiza* (Araceae) were lower than those of cooccurring diploids (Anneberg et al. 2023); however, a tetraploid fitness advantage was identified in saline environments (Bafort et al., 2023). Despite the competitive disadvantage of *S. polyrhiza* tetraploids compared to diploids in most treatments, tetraploids were found to invest more than their diploid progenitors into future growth through dormant propagules (Anneberg et al., 2023), which minimize the effect of variable conditions and are known to promote coexistence (Armstrong and McGehee 1980; Yuan and Chesson 2015). Investment in progagules and future growth by tetraploids could give them an advantage under large population fluctuations, a prediction that can be quantified in our model and would support tetraploid persistence in unstable environments.

Our model allows niche divergence among cytotypes to be driven by the environment while allowing independent, yet equal, survival. Coupled with stochasticity, niche independence may allow the persistence of one cytotype; however, it is unlikely that this interaction alone could drive the observed competitive advantage seen for tetraploids. Additional differentiation in adaptive ability between tetraploids and diploids, known to occur in nature, may influence the competitive advantage of the cytotypes during environmental stress. Across angiosperm phylogeny, duplicate gene retention following WGD has been linked to stress-related processes and may suggest that even paleopolyploids experience an adaptive advantage (Van de Peer et al. 2021; Wu et al. 2019). Similar adaptations have been confirmed in neopolyploids. For example, tetraploids have been identified to persist in high-salinity (Bafort et al. 2023; Chao et al. 2013; Scarrow et al. 2022; Thomas et al. 2022; Xu et al. 2021) and water-limited (Laport et al. 2017; Yang et al. 2014) environments. Despite common adaptations found in multiple systems, the mechanisms driving these responses are complex and difficult to model. For example, the interaction among ploidy, environment, and genotype has been found to influence cytotype-specific competitive advantage (Bafort et al. 2023; Van Drunen and Husband 2018).

### Concluding Remarks

Through our modeling approach, we found that long-run coexistence in mixed-ploidy populations is stabilized by increased selfing rates and pronounced reproductive isolation. Although neoautotetraploids initially face a numerical disadvantage, their ultimate fate is modulated by environmental stochasticity and can be quite variable. Under stressful environments, modeled here with an environmental noise regime, the dynamics among diploids, triploids, and tetraploids become much more complex. Our stochastic modeling approach revealed, for instance, that tetraploids have a competitive advantage over their diploid progenitors when environmental variance is high.

Developing stochastic population dynamic models linked to our current empirical understanding of mixed-cytotype populations is a research endeavor still in its infancy. While these modeling efforts improve our understanding of coexistence in mixed-cytotype populations, expanding their complexity is not simply a matter of bringing to bear additional computational effort. The promise of computational power may lure biologists to build models of complex dynamics, but many mathematical and programming decisions have to be made. Much mathematical and probabilistic study is necessary to ensure that the resulting predictions and explanations are not a by-product of, for instance, arbitrary computational decisions (see Ferguson and Ponciano 2014). One should proceed with further investigations once the results are fully understood. Taking this approach led us here to the surprising finding of the importance that gamete “stealing” has for tetraploid persistence in stressful environments.

Our modeling approach should be expanded to investigate the impact of additional processes on coexistence, such as temporally fluctuating unreduced gamete production, implicit niche competition, and variance in offspring success, to name a few. Of particular interest to us is completing our simulation framework by including genotype dynamics.

## Acknowledgments

Funding was provided to MLG from an NSF Graduate Research Fellowship (DGE-1842473). We thank R. G. Laport for feedback.

## Data and Code Accessibility Statement

This simulation framework into R software that is available via our package AutoPop: https://github.com/mgaynor1/AutoPop. Scripts associated with model evaluation and generation can be found in DRYAD:

## Notes

### Competing Interest Statement

The authors have declared no competing interest.

