## Supplemental for "Dynamics of mixed-ploidy populations under demographic and environmental stochasticities"

Michelle L. Gaynor[1,*]

Nicholas Kortessis[1,2]

Douglas E. Soltis[1]

Pamela S. Soltis[1]

José Miguel Ponciano[1]

1. University of Florida, Gainesville, Florida, 32611 2. Wake Forest University, Winston-Salem, North Carolina, 27109

### Supplementary Figures and Tables

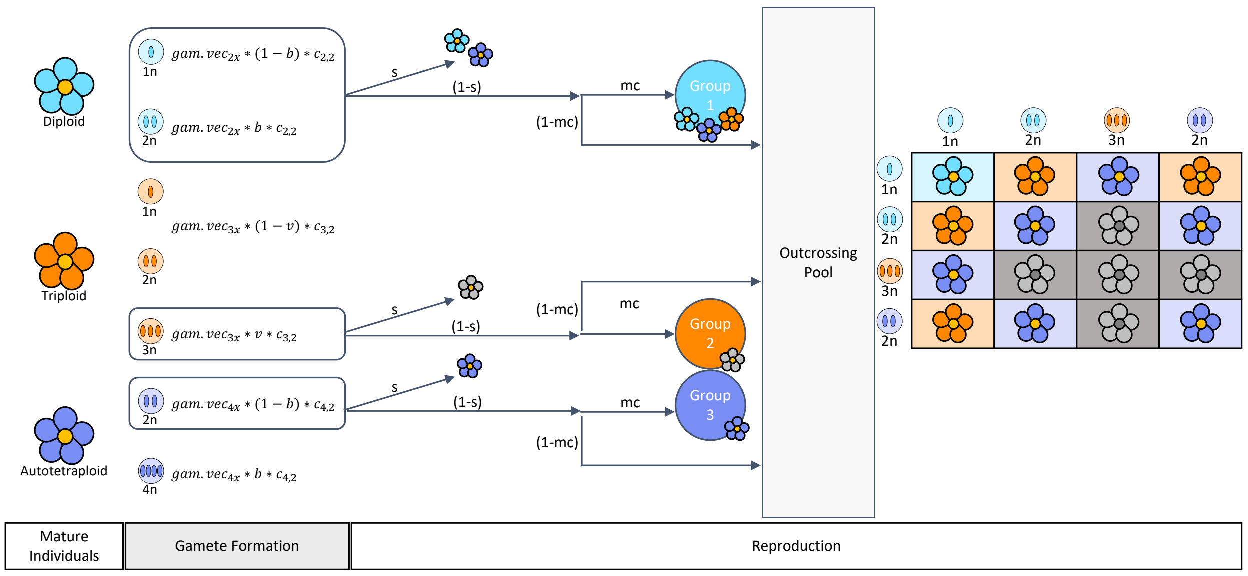


Figure S1: Visual display of reproduction including gamete formation ($gam.vec$, $b$, and $v$) and the union of gametes via sexual or asexual reproduction ($s$ and $mc$). The number of gametes of each type is calculated based on the number of mature individuals of each cytotype ($c_{2,2}, c_{3,2}, c_{4,2}$), the number of gametes produced by each individual ($gam.vec$), the frequency of unreduced gamete formation ($b$), and the proportion triploid gametes that will be viable ($v$). After gamete formation, selfing will occur based on a defined selfing rate ($s$), the remaining gametes will outcross. Of these outcrossing gametes, a proportion will only pair with gametes produced by like-cytotypes indicated by mating choice ($mc$), the remainder will freely pair in a outcrossing pool.

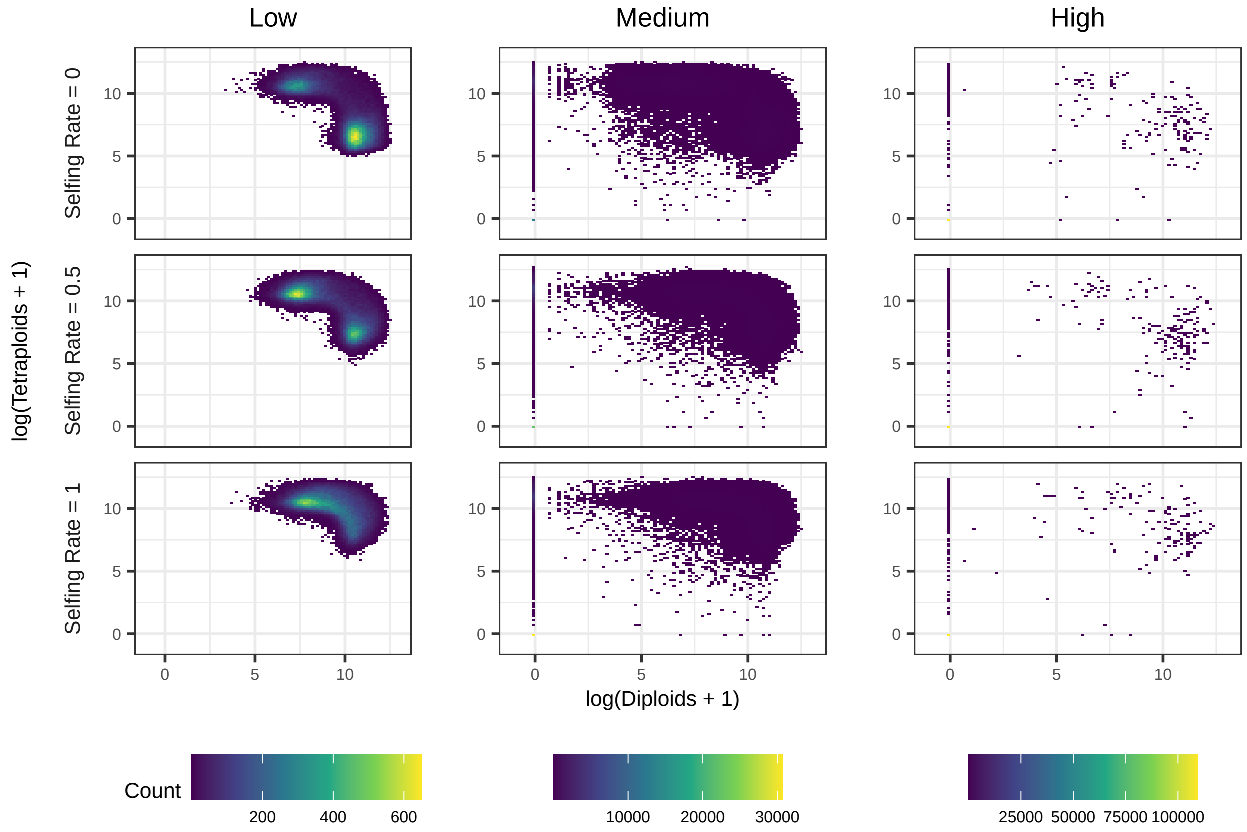


Figure S2: Density of diploids and tetraploids after burn-in for low, medium, and high environmental variance when the inital population size of 100. For low environmental variance we find two basins of attraction, where one cytotype is at high density and the other is at low density. For both medium and high environmental variance we observe a shift in the basin of attraction to extinction.

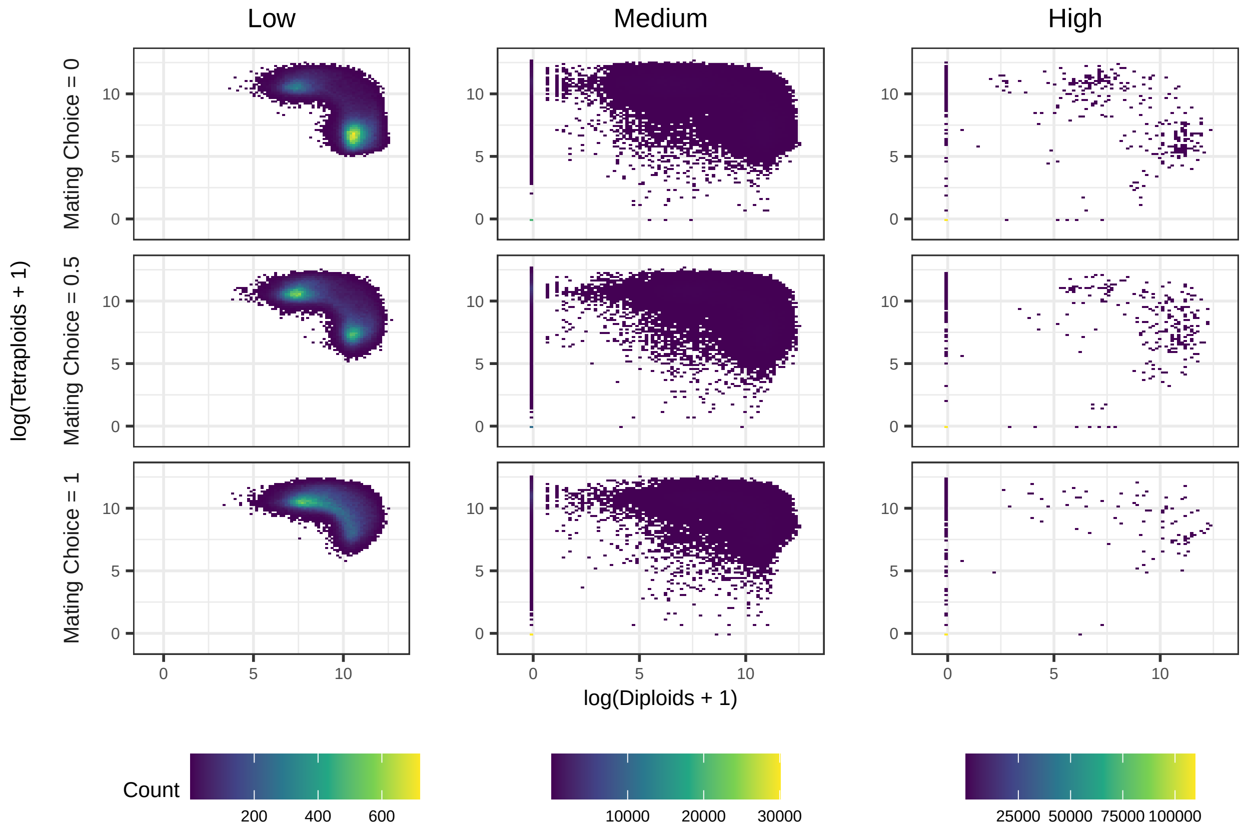


Figure S3: Density of diploids and tetraploids after burn-in for low, medium, and high environmental variance when the initial population size of 100. For low environmental variance we find two basins of attraction, where one cytotype is at high density and the other is at low density. For both medium and high environmental variance we observe a shift in the basin of attraction to extinction.

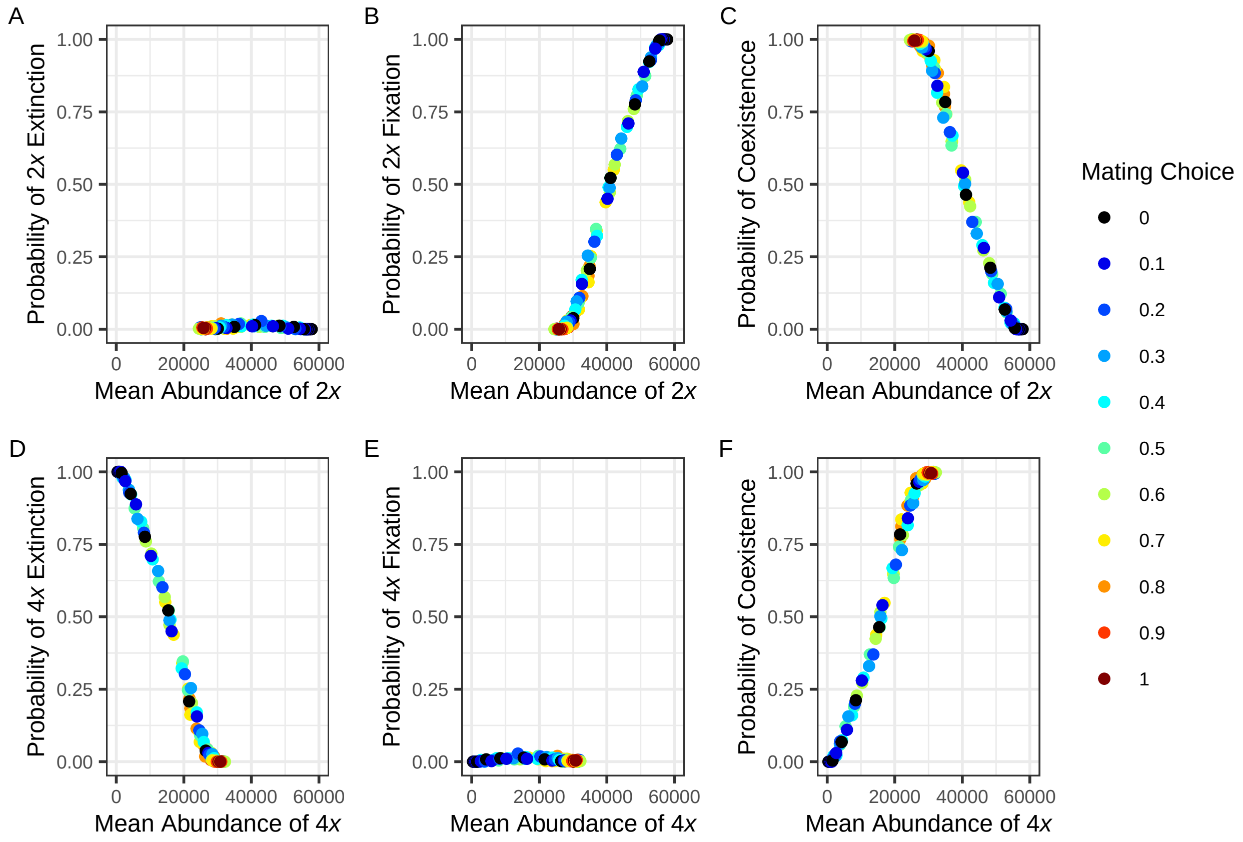


Figure S4: Probability of extinction (A, D), fixation (B, E), and coexistence (C, F) as function of the mean abundance of each cytotype for 500 generations after reaching stationarity. For tetraploids, increased mean abundance is observed with decreased probability of extinction. For diploids, we observe increased mean abundance after stationarity with increased probability of fixation. Mating choice shows no obvious relationship with the probability of extinction, fixation, or coexistence.

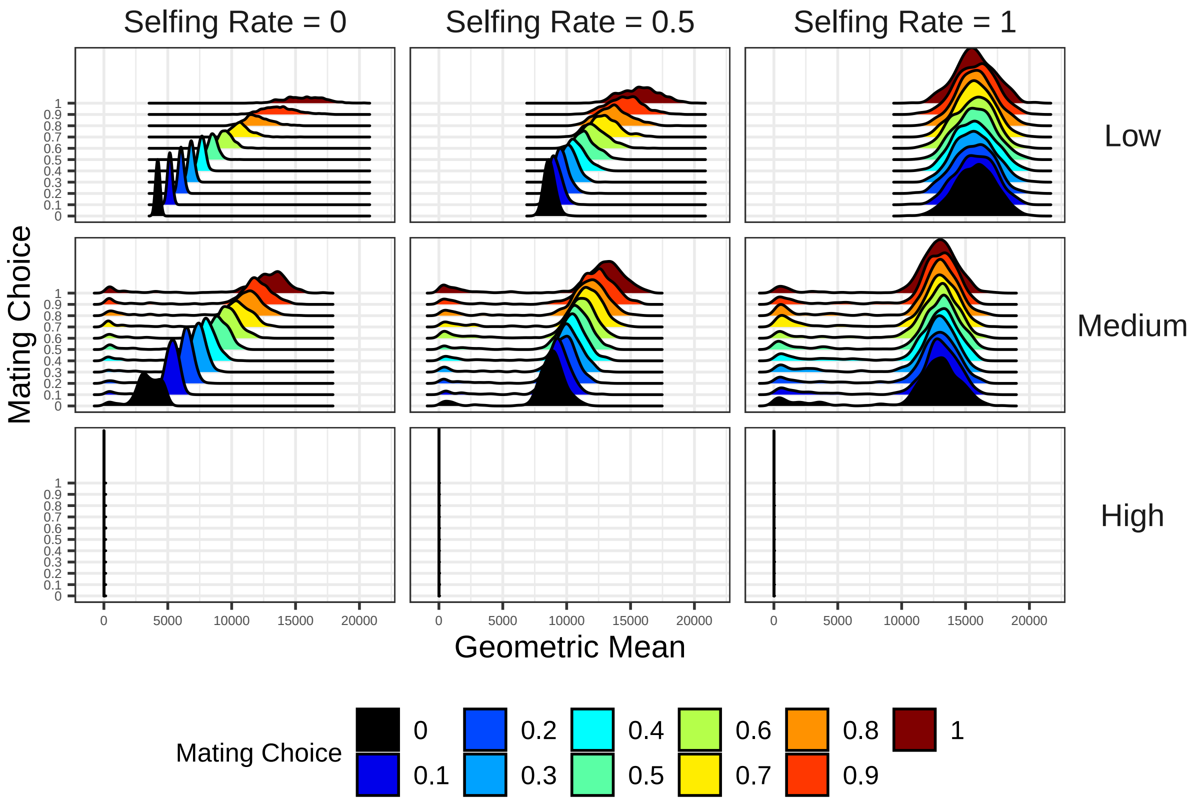


Figure S5: Long-run geometric mean for all 500 replicates for each model parameter combinations. For each simulated time series within each batch of 500 simulation runs, we computed the geometric mean over 500 generations after burn in of the abundances of diploids and tetraploids. Then we computed the geometric mean of these two geometric means. We then repeated the same calculation for each one of the 500 simulation runs. The distribution of these geometric means of the geometric means are here displayed as densities, for each parameter combination. Based on the resulting means, coexistence is defined as any value greater 0. At high environmental variance, we observe catastrophic stochasticity, as defined by Dennis et al. (1991).

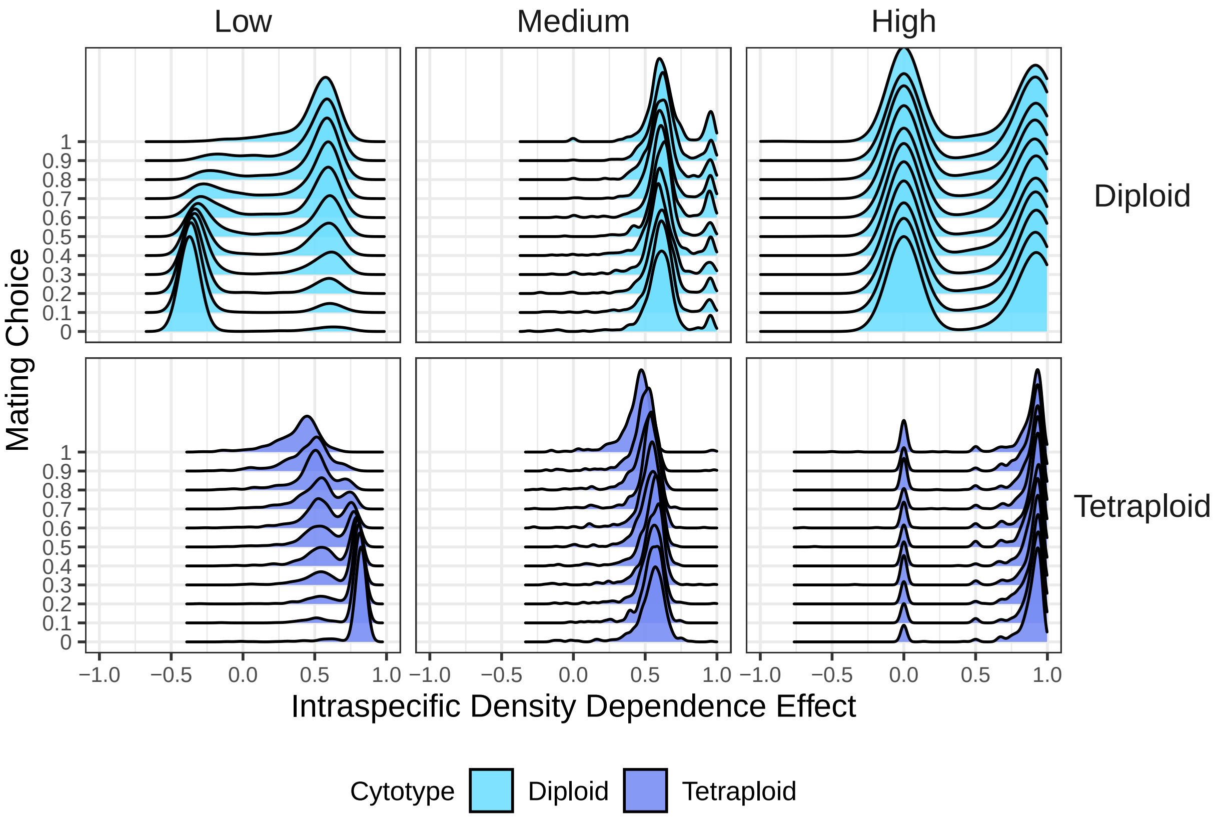


Figure S6: Intraspecific density dependence effect, as derived based on conditional least squares, for low, medium, and high environmental variance where the initial population size is equal to 100 and selfing rate is equal to 0.5. Values in the interval (-0.5, 0.5) denote strong density dependence, whereas values approaching 1 denote weak density dependence. Values of 1 denote in turn stochastic density independent growth. At low environmental variance, asymmetry in the coefficients between the diploid and tetraploid is more pronounced as selfing rate and mating choice decreases. At high environmental variance, tetraploids face weaker density dependence effect compared to diploids.

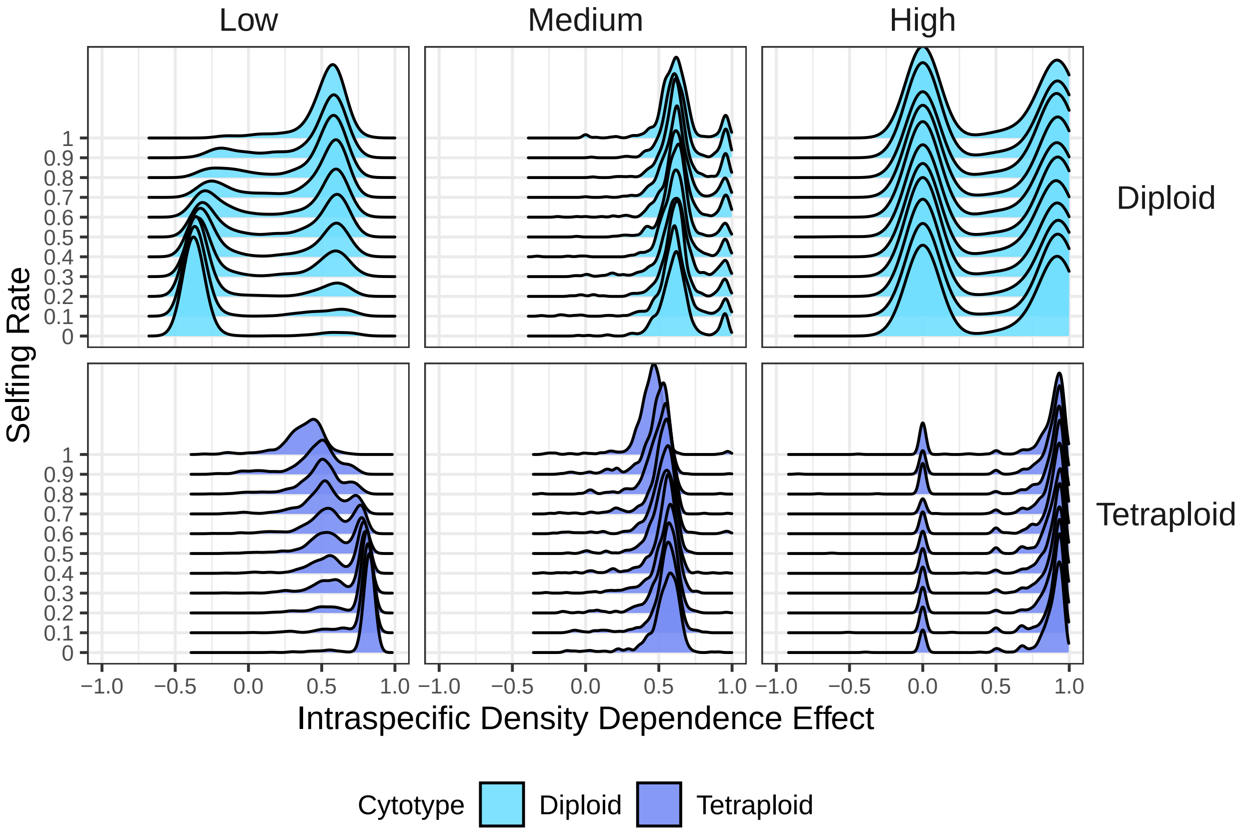


Figure S7: Intraspecific density dependence effect, as derived based on conditional least squares, for low, medium, and high environmental variance where the initial population size of 100 and selfing rate is equal to 0.5. Values in the interval (-0.5, 0.5) denote strong density dependence, whereas values approaching 1 denote weak density dependence. Values of 1 denote in turn stochastic density independent growth. At low environmental variance, asymmetry in the coefficients between the diploid and tetraploid are more pronounced as selfing rate decreases. At high environmental variance, tetraploids face weaker density dependence effect compared to diploids.
